# Broad-Spectrum Antiviral Efficacy of 7-Deaza-7-Fluoro-2’-C-Methyladenosine Against Multiple Coronaviruses *In Vitro* and *In Vivo*

**DOI:** 10.1101/2025.01.02.631052

**Authors:** Ian de Meira Chaves, Filipe Resende Oliveira de Souza, Celso Martins Queiroz-Junior, Leonardo Camilo de Oliveira, Ana Cláudia Dos Santos Pereira Andrade, Victor Rodrigues de Melo Costa, Danielle Cunha Teixeira, Felipe Rocha da Silva Santos, Talita Cristina Martins da Fonseca, Larisse de Souza Barbosa Lacerda, Rafaela das Dores Pereira, Jordane Clarisse Pimenta, Franck Amblard, Raymond F. Schinazi, Mauro Martins Teixeira, Vivian Vasconcelos Costa

**Author notes:** Both are corresponding authors and.

## Abstract

The Coronaviridae family has been implicated in several major epidemics over the past two decades, including those caused by SARS-CoV, MERS-CoV, and, most recently, SARS-CoV-2. The COVID-19 pandemic, driven by SARS-CoV-2, has led to over seven million deaths worldwide and has been associated with prolonged symptoms, chronic sequelae, and substantial socioeconomic disruptions. The limited availability of effective antiviral treatments, coupled with the ability of coronaviruses to mutate and evade immune defenses, underscores the urgent need for innovative antiviral agents. This study explores the efficacy of the nucleoside analogue DFMA as a potential antiviral agent against multiple Coronaviridae family members, including SARS-CoV-2 and two strains of murine hepatitis viruses (MHV-3 and MHV-A59). In vitro analyses demonstrated that DFMA effectively reduced the viral load in the supernatant of infected cells and enhanced cell viability for both MHV-3 and MHV-A59. Against SARS-CoV-2, DFMA showed a significant reduction in viral load, with a calculated Selectivity Index (SI) of 6.2. In vivo investigations further confirmed the antiviral potential of DFMA. In three distinct murine models—a severe COVID-19 model using MHV-3, a mild COVID-19 model employing MHV-A59, and a transgenic K18-hACE2 mouse model infected with SARS-CoV-2—DFMA administration significantly reduced viral loads in the lungs of infected mice. Additionally, DFMA mitigated inflammatory responses in all models by lowering levels of key inflammatory mediators, such as CXCL1, CCL2, and IL-6. These findings suggest that DFMA possesses broad-spectrum antiviral activity against coronaviruses and may serve as a promising therapeutic candidate for current and future coronavirus outbreaks. Further research is warranted to elucidate its mechanism of action and evaluate its efficacy in clinical settings.

**Importance:** Coronaviruses have caused significant outbreaks over the past two decades. Since 2020, COVID-19 has resulted in millions of deaths and lasting global impacts. The limited availability of effective antivirals and the virus’s ability to mutate and evade vaccines and monoclonal antibody therapy emphasize the urgent need for new treatments. This study investigates DFMA, a promising antiviral candidate, targeting SARS-CoV-2 and two related coronaviruses.Our promising results demonstrated significant antiviral activity of DFMA, not only against SARS-CoV-2 but also against other similar coronaviruses, indicating potential future use against COVID-19 and other possible coronavirus-related diseases.

## 1. Introduction

COVID-19 has caused over seven million deaths globally and has overwhelmed healthcare systems. Additionally, Sars-CoV-2 can lead to prolonged symptoms, known as Long COVID, which affects more than 100 million people worldwide (1–6) and is associated with significant socioeconomic impacts (7). Severe COVID-19 is associated with high viral titers commonly accompanied by an exacerbated host inflammatory response, characterized by massive production of inflammatory mediators known as cytokine storm (8, 9).

Currently, there are several vaccines approved for emergency use against COVID-19 and various vaccines are under development around the world. Meanwhile, a factor of concern was the speed at which new variants of SARS-CoV-2 emerged. Several variants have been described since the emergence of COVID-19, and mutations presented by new variants are constantly being associated with greater transmissibility and enhanced immune evasion in relation to vaccines in use and previous infections (10–14). For other human coronaviruses, a positive correlation between the severity of the disease and high viral titers in these individuals has already been demonstrated (15) and was recently reported in COVID-19 patients (16).

Given that the Coronaviridae family has repeatedly demonstrated its ability to cause significant public health crises, and with its viruses exhibiting high mutation rates, there is a critical need for effective treatments. Due to the short therapeutic window of the disease, it is imperative to identify new molecules capable of reducing viral titers during the acute phase, thereby improving both the immediate and long-term prognosis of patients.

Only three antiviral medications have already been approved by the US Food and Drug Administration (FDA) for the treatment of COVID-19. They are: i) Nirmatrelvir as a 3CL protease inhibitor (17). ii) Remdesivir (Veklury) (18, 19) and iii) Molnupiravir (Lagevrio) (20–22), both nucleoside analogs. Considering the scarcity of antiviral molecules, the search for alternative therapeutic options aimed at containing and/or alleviating cases of COVID-19, such as the use of antiviral drugs or monoclonal antibody therapies becomes necessary.

As a screening platform to assist in testing potential drugs, the murine coronavirus MHV (Mouse Hepatitis Virus) has been utilized. MHV, like SARS-CoV-2, is a betacoronavirus; however, it is a natural pathogen in mice capable of causing hepatitis and potentially leading to lethality in these animals (23). Working with the MHV requires biosafety level 2 laboratories and animal facilities, unlike the SARS-CoV-2, which necessitates a biosafety level 3 laboratory. Our group has demonstrated the usability of an infection model with the MHV-3 capable of causing severe pulmonary damage when inoculated intranasally. This model has been employed to study alternative methods of attenuation of the disease and to understand parameters related to viral replication and coronavirus acute pathology (24, 25). Additionally, MHV-A59 has been recently employed to study the acute lung infection as well as the long-term-induced neuropsiquiatric sequalae (26).

In this study, we propose to test the potential antiviral activity of 7-deaza-7-fluoro-2′-C-methyladenosine (DFMA) against different coronaviruses, including MHV and SARS-CoV-2. DFMA is a nucleoside analog that has potent antiviral activity and reduces the neuroinflammation induced by ZIKV infection *in vivo* (28), as well as DENV and WNV (29) and YFV (unpublished data). Therefore, in the current study, we evaluated DFMA efficacy both *in vitro* and *in vivo* using different models of coronavirus infection, including MHV and SARS-CoV-2. Our results indicate that DFMA is a potent antiviral capable of exerting activity against three members of the Coronaviridae family, including SARS-CoV-2. Therefore, we believe this preclinical study could serve as an initial stage in the development of this promising nucleoside analog.

## 2. Material and Methods

### 2.1 Viruses and Antiviral agent (DFMA)

The murine hepatitis virus strain MHV-3 (GenBank accession no. MW620427.1) and MHV-A59 (ATCC VR-764^TM^ available at https://www.atcc.org/products/all/VR-764.aspx) were acquired and propagated in L929 cells (ATCC, CCL1) for the generation of viral stocks. SARS-CoV-2 (Wuhan Hu-1 sample SP02BRA GenBank access number MT350282) was propagated in Vero CCL81 cells and was used in experiments conducted under BSL3 conditions. DFMA was synthesized in the Schinazi lab at Emory University as previously reported (29) and was at least 98% pure as determined by NMR and LC-MS-MS.

### 2.2 *In vitro* experimentation

L929 cells were cultured using DMEM medium, and Vero cells (BCRJ 0245) were cultured using RPMI medium. In both cases, the media were supplemented with 7% fetal bovine serum (FBS) and antibiotics for propagation and 2% fetal bovine serum for plating and during the assays. CALU-3 lung epithelial cells (BCRJ 0264 http://bcrj.org.br/celula/0264) were cultured in DMEM medium supplemented with 20% fetal bovine serum (FBS), 1% of non-essential amino acids, 2 mM de L-glutamine, 1 mM of sodium pyruvate, as well as antibiotics. All cells were maintained in an incubator at 37 °C and 5% CO_2_.

In both MHV-3 and SARS-CoV-2 infections, DFMA was tested in a 96-well plate being inoculated together with the virus at an MOI of 0.01 at concentrations of 3 μM, 10 μM or 30 μM. *In vitro* tests for the MHV were conducted using L929 cells, which were plated at 5 x 10^4^ cells per well and incubated for 24 hr at 37°C at 5% CO_2_. Medium was removed, and MHV-3 or MHV-A59 was added at MOI 0.01 along with different concentrations of DFMA. After 16 h, LDH assays were performed, and the supernatant was collected for viral load analysis.

*In vitro* SARS-CoV-2 experiments were conducted in partnership with the UFMG BSL3 facilities and the BSL4 of LANAGRO Laboratory, part of the Federal Network of Agricultural Defense Laboratories. Calu-3 were plated at 2,5 x 10^4^ cells per well and incubated under the same parameters described above. After 48 h, cells were infected with SARS-CoV-2 at MOI 0.01 along with 3 μM, 10 μM and 30 μM of DFMA. After an additional 24 h, DFMA was re-added at the same concentrations. Analyses were conducted 48 h post-viral inoculation. The supernatant was collected and stored at −80°C for viral titer assessment. Toxicity tests of DFMA on Calu-3 cells were performed in a BSL-2 environment, where cells were plated and incubated at the same conditions described above without infection, *i.e.* DFMA was added at different concentrations and re-added after more 24 h. After 48 h from the initial addition of DFMA, MTT assay was conducted.

#### 4.2.1 Plaque assay

Vero or L929 monolayers were seeded into 24-well plates and incubated for 24 hr at 37 °C with 5% CO_2_. Monolayers were then exposed to different dilutions of supernatants of *in vitro* tests or diluted organ samples for 1 h at 37 °C. Fresh semi-solid medium containing 1.2% carboxymethylcellulose (CMC) was then added, and the cultures were maintained at 37 °C for 72 h. After incubation, cells were fixed with a 10% formaldehyde solution for 60 min at room temperature and subsequently stained with 0.5% crystal violet for 30 min. Titer was expressed as pfu/mL.

### 4.3 MTT (3-(4,5-dimethylthiazol-2-yl)-2,5-diphenyltetrazolium bromide)

The MTT method was employed to assess cell viability, using a concentration of 0.5 mg/mL diluted in DMEM medium. The solution was applied to the cells and then incubated at 37 °C with 5% CO_2_ for 2 hr. After incubation, the content was removed and the salt formed by the MTT reaction was dissolved by adding 100 µL of DMSO. Absorbance was measured using a plate spectrophotometer at OD of 540 nm.

### 4.4 LDH (lactate dehydrogenase)

For the LDH assay, a kinetic kit from Bioclin (catalog no. K014, Bioclin, Belo Horizonte, Brazil) was used, following the methodology recommended by the manufacturer with modifications. Test was conducted in 96-well plates, with 4 µL of cellular supernatant added. Subsequently, 200 µL of the kit’s working solution was added, and the absorbance was measured immediately at 340 nm. Three readings were taken with a one-minute interval between readings to calculate substrate consumption per minute. The values were expressed in absorbance (Δ/min).

### 4.5 *In vivo* infections and DFMA treatment

C57BL/6 mice aged 6 to 8 weeks, supplied by the Central Animal Facility of the Federal University of Minas Gerais (UFMG) were supplied by Biotério Central – UFMG and used for the experiments involving MHV. K18-Human ACE2 Transgenic mice originally obtained from The Jackson laboratories (https://www.jax.org/strain/034860), created and maintained at Departamento de Bioquímica e Imunologia from UFMG were used for SARS-CoV-2 experiments. In both instances, for intranasal infection, mice were anesthetized subcutaneously with a solution containing 80 mg/kg of ketamine and 15 mg/kg of xylazine. Mice infected with MHV or SARS-CoV-2 received 30μL of the viral inoculum containing 1 x 10^3^ PFU for MHV-3 as described by Andrade et al (24), 1 x 10^4^ PFU for MHV-A59 (26) and 2 x 10^4^ for SARS-CoV-2 (27). This study was previously submitted to the Ethics Committee on Animal Experimentation at UFMG and approved under protocols 338/2023, 140/2023 (CEUA-UFMG) and 191/2021 (CEUA-UFMG).

For evaluation of the effect of DFMA during viral infection, mice were treated BID with 10 mg/kg of DFMA, starting the treatment two days before infection (28). Mice were randomly divided into groups, weighed, and monitored daily. After infection, mice that reached 80% of their initial body weight were euthanized in accordance with the rules of the Ethics Committee for Animal Experimentation (CEUA-UFMG). For analysis, mice were euthanized on the peak of viral replication or lung damage, represented as day 3 after MHV-3 infection (24), second day after MHV-A59 infection, and third day after SARS-CoV-2 infection. Analysis, included inflammatory scoring in the lungs, quantification of chemokines and cytokines in the plasma and lungs of the animals, viral load quantification by qPCR in the lungs, immunohistochemistry for visualization of Spike protein of SARS-CoV-2 or MHV and hematological analyses.

### 4.8 Viral quantification by qPCR

The viral RNA from the tissue was extracted using the viral RNA extraction kit QIAamp®□ Viral RNA following the manufacturer’s instructions, with adaptations, where the tissues were removed from RNAlater and homogenized in the lysis buffer of the kit. Samples were eluted and quantified on a spectrophotometer (NanoDrop™□, Thermo Scientific). cDNA production was performed using 500 ng of RNA, using the kit iScript™□ gDNA Clear cDNA Synthesis Kit (BIO-RAD), following the manufacturer’s instructions. The obtained cDNA was diluted 10x for use in the qPCR reaction. In the reaction, Fast SYBR™□ Green Master Mix (Applied Biosystems™□) was used with specific primers for MHV at concentration of 5 nM: Forward 5′-CAGATCCTTGATGATGGCGTAGT-3′; Reverse 5′-AGAGTGTCCTATCCCGACTTTCTC-3′. A standard curve was obtained through the RNA extraction from a known PFU quantity of MHV and the results were expressed in arbitrary units/500ng of cDNA. qPCR for detecting SARS-CoV-2 was performed using 2019-nCoV RUO kit (IDT) for N1 region and reaction ran in CFX Opus 96 (BIO-RAD). Standard was produced from a known number of copies by 2019-nCoV_N Positive Control (IDT). The results were expressed by number of copies/ 50ng of RNA.

### 4.9 Hematological analyses

Blood collection was performed after euthanasia and assessment of parameters on the veterinary hematological analyzer for the experiments with MHV-3 and MHV-A59 in BSL-2 laboratory (MEK 6550J/K – Celltak). Blood was then centrifuged at 4°C at 5,000 RPM for 10 min, and the supernatant was collected and frozen at −80°C for subsequent analysis.

### 4.10 Histopathological analyses

Histopathological analysis was performed in the lung of mice which were collected and fixed in 4% buffered formalin solution after euthanasia. Samples were then embedded in paraffin, sectioned using a microtome, mounted on histological slides, processed and stained with hematoxylin and eosin. Analyses were carried out using a system previously described by Andrade et al. (2021) (24), performed by a blinded pathologist.

### 4.11 Immunohistochemistry (IHC)

Histological lung sections from infected animals treated or untreated with DFMA were processed for spike protein staining using immunohistochemistry. The process involved a streptavidin-biotin protocol. Endogenous peroxidase was blocked using 0.3% hydrogen peroxide, and antigen retrieval was performed with 1.266 mM EDTA buffer at pH 8. Sections were incubated with primary Anti-MHV nucleocapsid NR-45106 - 7003428 provided by BEI resources at 1:100 dilution for MHV staining. For SARS-CoV-2 staining, SARS-CoV-2 (2019-nCoV) Spike Antibody (Sino biological - 40591-T62) at a 1:500 dilution was used. Subsequently, the section was incubated with the secondary Mouse on Mouse kit purchased from Vector Labs (USA) for MHV staining and the anti-rabbit secondary ABC kit (Vector laboratories) for SARS-CoV-2 and both were developed with DAB chromogen counterstained with hematoxylin. Negative reaction controls were performed on sections without primary antibodies added.

### 4.12 ELISA

Quantification of the chemokines CXCL1, CCL2, and the cytokine IL-6 were performed using the mouse DuoSet enzyme-limited immunosorbent assay (ELISA) system (R&D Systems), as described by Andrade et al (2021) (24).

### 4.13 Statistical analyses

Data were expressed as mean ± standard error of mean (SEM) or just as mean. The normality test was applied using the Shapiro-Wilk test. Means were compared using analysis of variance by one-way ANOVA, followed by Tukey test for normal distribution and Kruskal-Wallis plus Dunn’s test for sampling without normal distribution and for analyses involving only two groups, the t-test was employed. Results were considered significant when the value of p < 0.05. For the construction of the graphs, GraphPad PRISM 9 software was used. The exclusion of outlier values was performed through the GraphPad PRISM software platform itself.

## 3. Results

### DFMA antiviral activity against MHV-3 and MHV-A59 *in vitro*

To assess the potential DFMA’s antiviral capability against coronaviruses, an *in vitro* evaluation was first conducted **(Fig. 1A)**, testing different concentrations of the drug within a pre-analyzed spectrum (28). First, MHV-3 was used, and *in vitro* tests indicated that MHV-3 induced the formation of numerous syncytia in L929 cells at an MOI of 0.01 **(Fig. 1B)**. DFMA at 30 μM partially restored the original morphology of L929 cells, as shown in representative images (**Fig. 1B**). DFMA exhibited notable antiviral activity, reducing viral titers in the supernatant of infected cells, with effects observable from 10 µM to 30 µM, with a 1,000-fold reduction in viral titers at 30 µM (**Fig. 1C**). Furthermore, cell death of the L929 monolayer was quantified using the LDH assay. MHV-3 induced a significant increase in cellular death, while DFMA, starting from 10 µM, partially reversed this phenotype. However, at a concentration of 30 µM, DFMA completely restored LDH levels comparable to uninfected cells (**Fig. 1D**).

**Figure 1.**
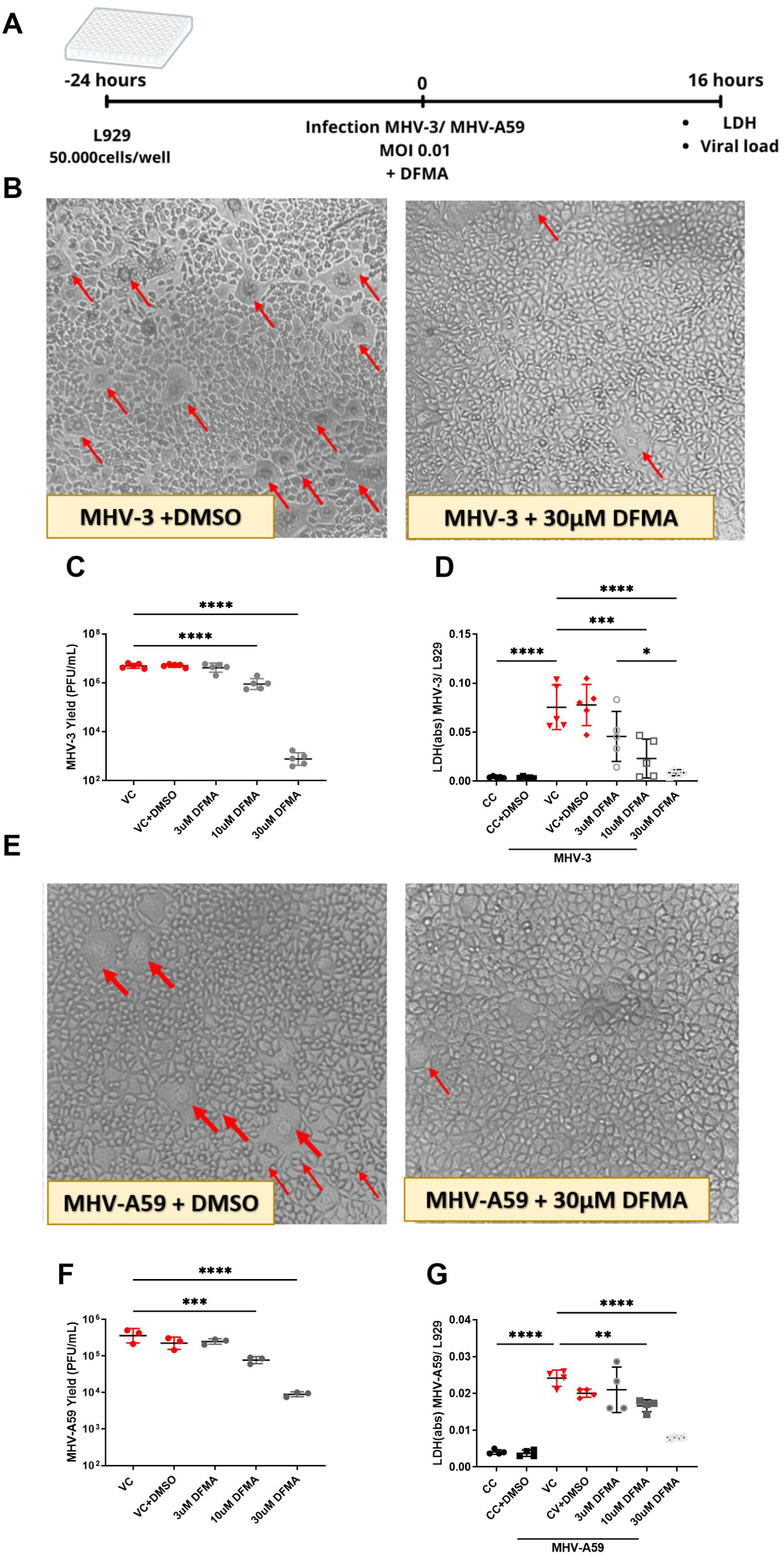
DFMA exhibits antiviral effects in vitro against MHV-3 and MHV-A59. L929 cells were infected with MHV at an MOI of 0.01, and different concentrations of DFMA (3, 10, and 30 μM) were added simultaneously with the viral inoculum. Cell controls (CC) received only DMEM plus 1% FBS; Cell Control + DMSO (CC+DMSO) received DMEM plus 1% DMSO; Viral Control (VC) received only viral content; and VC + DMSO received viral content plus 1% DMSO. The experimental design is shown in (A). After 16 hours, cytopathic effects with or without DFMA treatment were photographed, with syncytia indicated by arrows (B). The in vitro assay included viral titer analysis in the cell supernatant (C) and an LDH assay to assess cell death, expressed as Δ absorbance/min (D). The same parameters were evaluated for MHV-A59, including representative images of the L929 monolayer (E), viral titer in the supernatant (F), and LDH assay results (G). Statistical analysis was performed using one-way ANOVA followed by Tukey’s post-test. *P < 0.05; **P < 0.01; ***P < 0.001; ****P < 0.0001.

Further the *in vitro* antiviral potential of DFMA against the MHV-A59 was assessed. MHV-A59 also induced the formation of numerous syncytia in the L929 cell monolayer after 16 h post-inoculation. DFMA at 30 μM completely restored cell morphology (**Fig. 1E**). DFMA also showed a significantly reduced viral titer even at 10 µM **(Fig. 1F)**. At higher concentrations, DFMA achieved a reduction of over 10 times in the recovered viral titers from the supernatant at 30 μM when compared to non-treated cells **(Fig. 1F).** Cell death analysis demonstrated that, like MHV-3, MHV-A59 infection resulted in a significant increase in cell death **(Fig. 1G)**. DFMA prevented those parameters of cell death almost completely at 30 μM. **(Fig. 1G)**.

### DFMA treatment exhibits antiviral effects and improves disease parameters in a severe COVID model using MHV-3

MHV-3 has already been demonstrated as a pathogen capable of causing severe disease in mice, being considered a good model for severe acute COVID-like illness (24). Therefore, given that intranasal infection with the MHV-3 serves as a model for COVID, capable of mimicking severe COVID, the antiviral capacity of DFMA within the context of MHV-3 infection was evaluated *in vivo* **(Fig. 2A)**.

**Figure 2.**
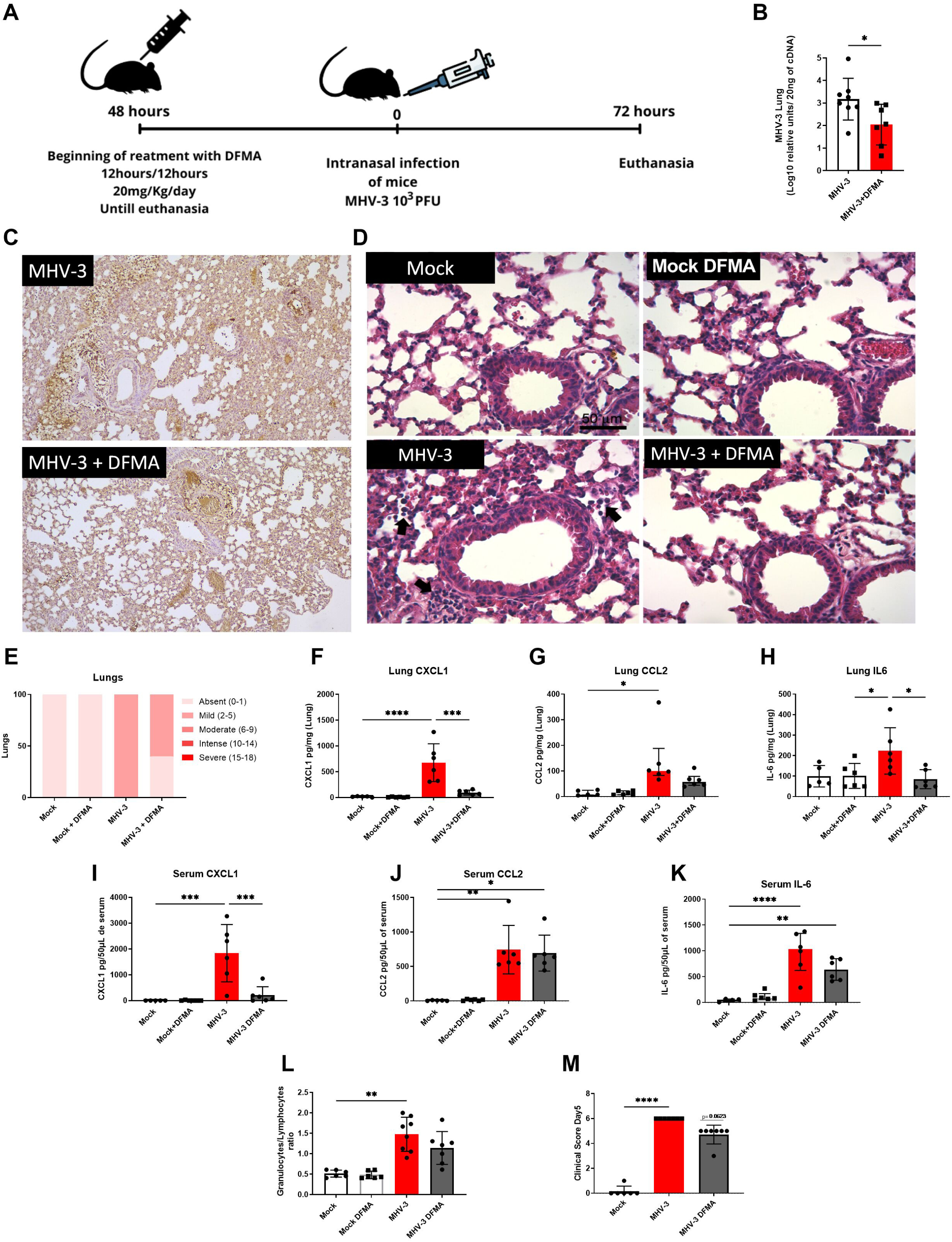
Effects of DFMA treatment against MHV-3 infection in mice. Mice were treated every 12 hours (B.I.D), starting two days before infection, with a total dose of 10 mg/kg, until the second day post-infection. The experimental design is illustrated in (A). Mice were infected with MHV-3 and euthanized on day 3 post-infection. A second experiment was conducted to assess clinical parameters on day 5 post-infection. Viral load in lung tissue was quantified by RT-qPCR and expressed as Relative Units/20 ng of cDNA (B). Immunohistochemical staining of MHV-3 nucleocapsid in the lungs are shown (C). Representative lung images from MHV-3-infected mice, with or without DFMA treatment, are shown. In untreated infected mice, increased inflammatory cell infiltration can be observed in lung tissue. Arrows indicate the presence of leukocytes in the alveolar and perivascular regions (D). Contingency representation of inflammatory score (E). Cytokine and chemokine quantifications in lung tissue are shown for CXCL1 (F), CCL2 (G), and IL-6 (H). The same mediators were also quantified in serum: CXCL1 (I), CCL2 (J), and IL-6 (K). Blood samples were collected, and the granulocyte-to-lymphocyte ratio was determined (L). The clinical score was evaluated on day 5 post-infection (M). Group differences were analyzed using one-way ANOVA followed by Tukey’s test for panels (E), (F), (G), (H) and (K); Kruskal-Wallis followed by Dunn’s test for panels (J), (L), and (M); and a t-test for two-group comparisons. *P < 0.05; **P < 0.01; ***P < 0.001; ****P < 0.0001. n = 6–8.

Mice infected with MHV-3 had the presence of viral RNA in their lungs at the third day after infection. Interestingly, treatment with DFMA led to a remarkable reduction of approximately 90% in viral load within the lungs of infected mice **(Fig. 2B)**.

The group of mice infected with MHV-A59, all the mice showed a mild inflammatory score, whereas in the group treated with DFMA, 40% of the mice presented absent inflammatory score **(Fig. 2C).** Histological analyses showed a higher inflammatory score in the lungs of mice infected with MHV-3, primarily characterized by a cellular infiltrate of polymorphonuclear cells in the perivascular area and in the alveolar space, along with hyperplasia of the alveolar walls **(Fig. 2D).** The inflammatory score induced by MHV-3 infection was only partially reduced with DFMA treatment, although the difference was not statistically significant **(Fig. 2E)**.

Recognizing the importance of the cytokine storm within the context of COVID-19, quantification of chemokines and cytokines present in the lung tissue was performed. MHV-3 infection induced an increase in the levels of the chemokines CXCL1 **(Fig. 2F)** and CCL2 **(Fig. 2G),** as well as an increased in the levels of the cytokine IL-6 **(Fig. 2H)**. Treatment with DFMA not only was able to reduce the viral load, but this reduction also reversed the increased levels of CXCL1 **(Fig. 2F)** and IL-6 **(Fig. 2H)** when compared to the vehicle-treated group.

Besides the lungs, MHV-3 was also able to systemically increase the levels of CXCL1 **(Fig. 2I)**, CCL2 **(Fig. 2J)**, and IL-6 **(Fig. 2K)** in the serum of infected animals. Interestingly, treatment with DFMA reversed this increase and restored the baseline levels of the chemokine CXCL1 **(Fig. 2L)**, although it did not show a significant difference for CCL2 **(Fig. 2J)** and IL-6 **(Fig. 2K)**. Systemically, granulocyte/lymphocyte ratio was evaluated. At the peak of the COVID pandemic, the neutrophil-lymphocyte ratio was considered an important marker for predicting patient prognosis (30,31). In our model, mice infected intranasally by MHV-3 and euthanized three days after showed increased granulocyte/lymphocyte ratio, but DFMA was not able to reverse this parameter (p = 0.197) **(Fig. 2L)**. Similarly, MHV-3 infection caused a significant increase in the clinical score of mice, and DFMA treatment failed to improve their condition (**Fig. 2M**).

### DFMA exhibited antiviral activity against MHV-A59 and improved inflammatory parameters *in vivo*

Given that COVID-19 complications can extend beyond the acute phase, the antiviral efficacy of DFMA was also evaluated in a mild COVID model using MHV-A59 **(Fig. 3A)**. MHV-A59 infection in mice was associated with the recovery of high viral RNA copies in the lungs, measured by RT-qPCR. Treatment with DFMA decreased the quantification of viral RNA copies **(Fig. 3B).** Representative images were also taken of lung histological sections showing a large inflammatory infiltrate in the lung tissues of untreated infected mice, which was not observed in mice treated with DFMA **(Fig. 3D)**. The group of mice infected with MHV-A59 showed a mild inflammatory score in 50% of the animals, whereas in the group treated with DFMA, only 14,3% presented alterations in the inflammatory score and reached the ‘’mild’’ level of score, while the remaining mice were classified as having an absent score **(Fig. 3E)**.

**Figure 3.**
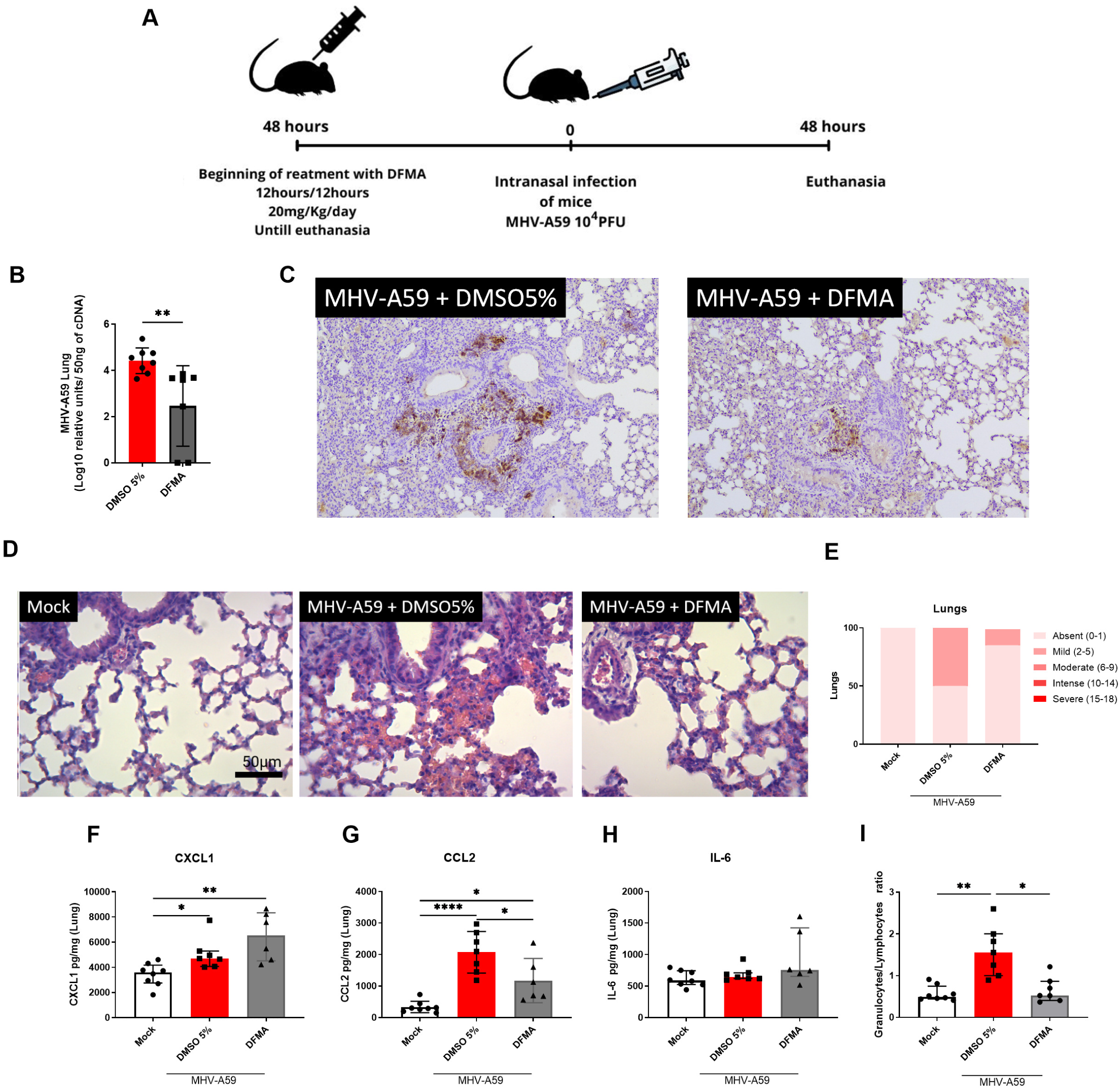
Effects of DFMA against MHV-A59 in mice. Mice were treated following the same protocol used previously for MHV-3. Mice were infected with MHV-A59 and divided into groups, with or without DFMA treatment. Two days post-infection, mice were euthanized and samples were collected. The experimental design is illustrated in (A). Viral load in the lungs was quantified by RT-qPCR and expressed as Relative Units/50 ng of cDNA (B). Immunohistochemical staining of MHV-A59 nucleocapsid in the lungs was shown in (C) in addition to representative histological analysis of lung tissue highlights inflammatory infiltrates and hemorrhage in untreated infected mice (D). Contingency representation of Inflammatory Score (E). Chemokine levels of CXCL1 (F) and CCL2 (G), along with the cytokine IL-6 (H), were measured in lung tissue samples. The granulocyte/lymphocyte ratio was assessed from blood samples (I). Group differences were evaluated by one-way ANOVA followed by Tukey’s test for panel (G), Kruskal-Wallis followed by Dunn’s test for panels (E), (F), (H), and (I), and a t-test for analyses involving only two groups. *P < 0.05; **P < 0.01; ***P < 0.001; ****P < 0.0001. n = 7–8.

In addition to assessing viral load and confirming that DFMA indeed exerted antiviral activity within the context of MHV-A59 infection, its ability to improve inflammatory parameters were also analyzed. MHV-A59 infection was associated with an increase in the levels of CXCL1 in both the lungs of infected mice treated and untreated with DFMA **(Fig. 2F)**. Upon evaluating the chemokine CCL2, a substantial elevation induced by MHV-A59 infection was detected in untreated mice. However, following DFMA treatment, mice exhibited diminished levels of this chemokine **(Fig. 2G)**. No difference between groups was detected when IL-6 was measured **(Fig. 2H)**. Additionally, MHV-A59 infection led to an increase in Granulocytes/Lymphocytes ratio **(Fig. 2I)**. Interestingly, in this model, treatment with DFMA decreased such ratio when compared to untreated mice **(Fig. 2H)**.

### DFMA exerted antiviral effects against SARS-CoV-2 *in vitro* and *in vivo*

To assess the potential antiviral effect of DFMA directly against the SARS-CoV-2 *in vitro*, immortalized lung epithelial cells (Calu-3) were infected at MOI 0.01 and treated with different DFMA concentrations **(Fig. 4A)**. DFMA at 30 µM induced a reduction of approximately 15-fold in the recovered viral titer when compared to the supernatant from untreated cells **(Fig. 4B)** with an EC_50_ value of 8.2 **(Fig. 4C).** To evaluate DFMA cytotoxicity in Calu-3 cells, cell monolayer was exposed to different DFMA concentrations, resulting in a CC_50_ of 51.0 **(Fig. 4C)** and an SI of 6.2, being considered a potent and promising antiviral candidate.

**Figure 4.**
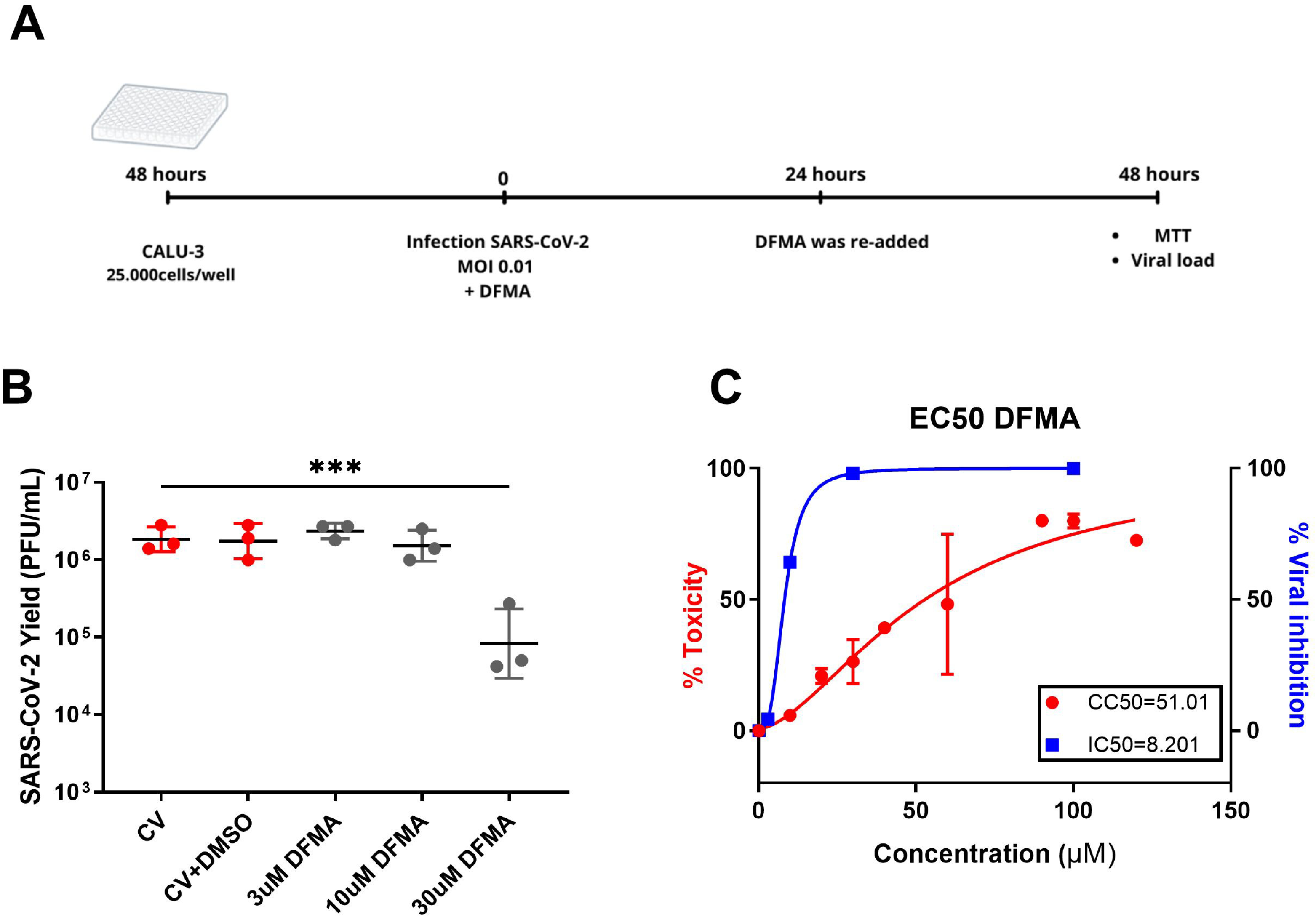
Antiviral activity of DFMA *In vitro* against SARS-CoV-2. Calu-3 cells were infected by SARS-CoV-2 at MOI = 0.01, along with DFMA added simultaneously in concentrations ranging from 3 to 30 μM. VC (Viral Control) received only virus, and VC+DMSO received virus plus 1% DMSO. After 24 hr exposure to DFMA, fresh solution of DFMA was applied to the corresponding wells. Experimental design was represented **(A).** A titration of the supernatant was performed **(B)**. One-way ANOVA followed by Tukey was performed ***P < 0.001. Additionally, a cytotoxicity assay was conducted by MTT to determine the CC_50_ of 51.0 for DFMA and the EC_50_ of 8.2, assessed by nonlinear regression. This allowed the calculation of the Selectivity Index SI = 6.2 of DFMA **(C)**.

An *in vivo* experiment was conducted using K18-Human ACE2 mice infected with SARS-CoV-2 and euthanized on the third day post-infection **(Fig. 5A)**. The infection resulted in a high recovery of viral RNA copies, and interestingly, DFMA treatment effectively reduced its levels in the lungs, as measured by qPCR **(Fig. 5B).** This reduction can be effectively detected with the staining of SARS-CoV-2 Spike protein in lung samples **(Fig. 5C)** validating all previous results and presenting DFMA as a potential antiviral for coronaviruses. Like in MHV infection, in addition to the viral load results, lung histological sections revealed a substantial inflammatory infiltrate in the lung tissues of untreated infected mice, which was less intense in most of the mice treated with DFMA **(Fig. 5D)**. The group of mice infected with SARS-CoV-2 showed a mild inflammatory score in 85% of the animals, whereas in the group treated with DFMA, only 12.5% presented alterations in the inflammatory score, with the remaining 87.5% considered to have absent score. **(Fig. 5E)**.

**Figure 5.**
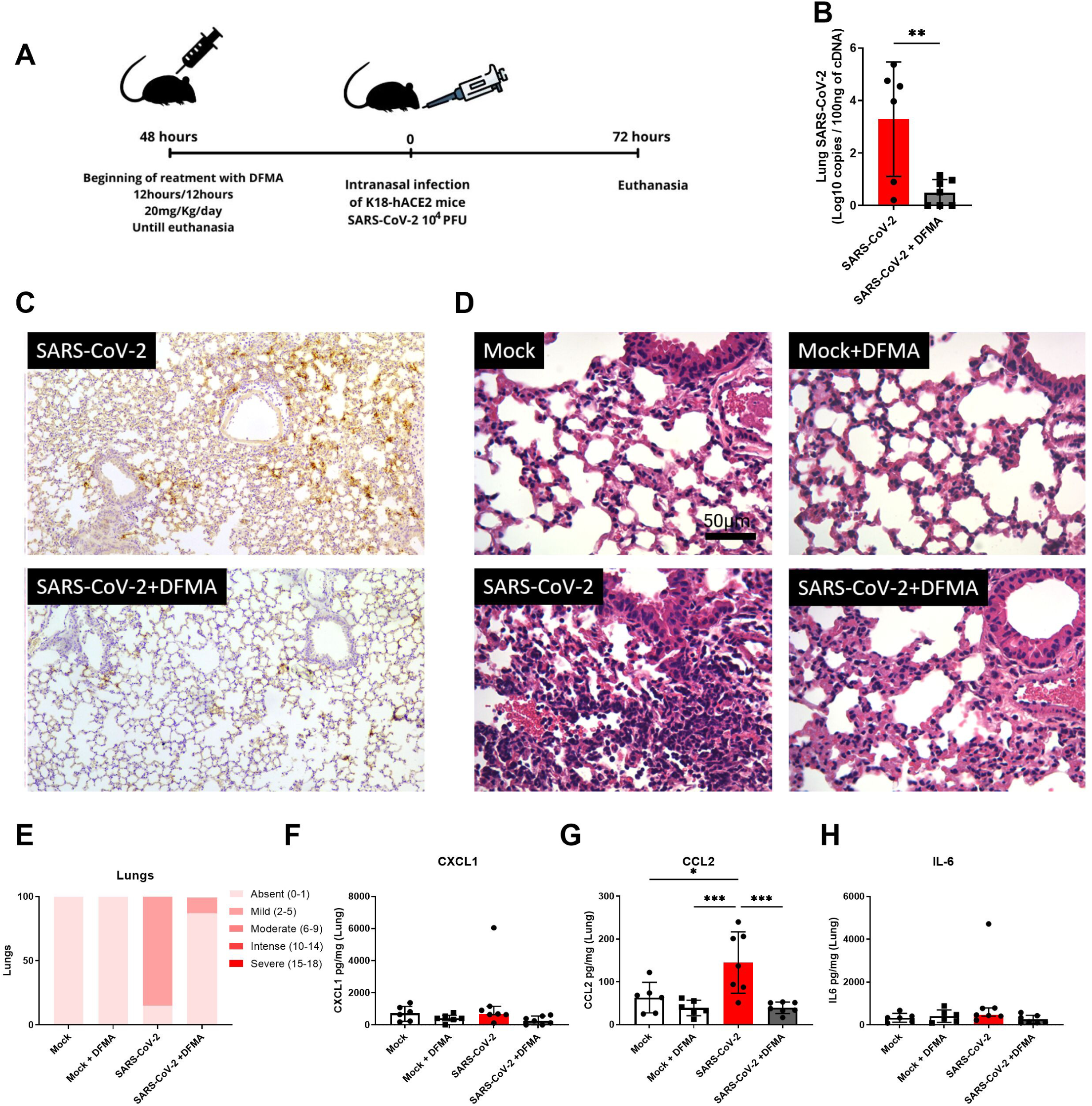
Effects of DFMA treatment on SARS-CoV-2 infection in K18-hACE2 mice. K18-hACE2 mice were treated with 10 mg/kg of DFMA twice daily (B.I.D) starting two days before infection and continuing until the second day post-infection. Mice were euthanized on the third-day post-infection with SARS-CoV-2. The experimental design is illustrated in (A). Viral load in the lungs was measured by quantifying viral RNA using RT-qPCR (B). Lung tissue was also collected for histological analysis, with immunohistochemical staining of the SARS-CoV-2 Spike protein (C). Representative histological images display inflammatory infiltrates in infected mice treated with vehicle control (D). Contingency Histogram of Inflammatory Score (E), alongside the quantification of inflammatory mediators, including CXCL1 (F), CCL2 (G), and IL-6 (H). Differences between groups were analyzed using one-way ANOVA followed by Tukey’s test for panel G, and Kruskal-Wallis followed by Dunn’s test for panels E, F, and H. For comparisons involving only two groups, a T-test was used. *P < 0.05; **P < 0.01; ***P < 0.001; ****P < 0.0001; n = 7-8.

Additionally, the quantification of inflammatory mediators in the lung tissues of mice infected with SARS-CoV-2 was also assessed. No differences were detected at 3 dpi in the lung levels of CXCL1 levels **(Fig 5F)** and IL-6 (Fig 5H). Conversely, analysis of the chemokine CCL2 revealed a significant increase in its levels in the lungs of SARS-CoV-2-infected animals. Notably, DFMA treatment effectively reduced CCL2 levels, restoring them to baseline **(Fig. 5G)**.

## 4. Discussion

*Coronaviridae* family includes various viruses capable of infecting humans, birds, rodents, and other animal species (32). Some coronaviruses are globally distributed and capable of causing mild respiratory infections in humans, such as 229E, OC43, NL63, and HKU1 (32). In addition to mild infections, coronaviruses have also been responsible for significant epidemics leading to serious respiratory complications over the past 20 years, such as severe acute respiratory syndrome (SARS) caused by SARS-CoV (WHO, 2003), resulting in profound economic, social, and health impacts (33); Middle East Respiratory Syndrome caused by MERS-CoV and more recently, in 2019, COVID-19 caused by the SARS-CoV-2 virus. In this study, the potential antiviral effects of a nucleoside analog, DFMA, were evaluated in various coronavirus infection models. Both *in vitro* and *in vivo* results demonstrated the promising antiviral activity of DFMA against MHV-3, MHV-A59, and SARS-CoV-2. In addition to reducing viral load across all three in vivo models, DFMA significantly decreased levels of key inflammatory mediators, including CXCL1, CCL2, and IL-6. This suggests that the reduction in viral load may also be modulating and mitigating the excessive inflammatory response, a hallmark of severe COVID-19 cases.

The COVID-19 has proven to be a disease of profound and enduring impact. In addition to the direct fatalities attributed to the illness and the consequential strain on healthcare systems worldwide, it exerts significant influence on various socio-economic dimensions (7). Notably, acute symptoms of COVID-19 are not the only concerning aspects of this syndrome. There is also the long COVID syndrome, a multisystemic and heterogeneous syndrome characterized by various prolonged systemic alterations that extend beyond the initially reported acute pulmonary complications. Studies demonstrate that the symptoms and consequences associated with SARS-CoV-2 infection can persist for months, potentially leading to irreversible and irreparable sequelae in many fields (2, 4, 5). Currently, there are several vaccines approved on an emergency basis that have played a significant role in reducing and mitigating the disease cases (34). A major cause for concern is the emergence of new variants carrying alterations, particularly related to the viral Spike protein that has been demonstrated to be associated with immune evasion and increased transmissibility (10–14, 35). COVID-19 pandemic once again highlights the need for new molecules capable of reducing viral replication or even interrupting the viral replication cycle.

Here, we demonstrated the antiviral activity of DFMA, a nucleoside analog, against coronavirus infection. In previous studies, it has been demonstrated that treatment with DFMA was able to reduce the number of viral RNA copies in the supernatant of cells infected with DENV-1, DENV-2, DENV-3, and DENV-4, as published by Zandi et al. (29). In this same study, *in vivo* analyses demonstrated DFMA’s ability to decrease viral titers of viable particles in the plasma, spleen, liver, and brain when administered at a dose of 10 mg/kg/day starting from the date of infection i mice deficient for type I interferon receptors (A129), as well as being associated with systemic improvements, such as reduced tissue damage, recovery of platelet levels, and lower quantifications of systemically released inflammatory mediators (29).

The promising antiviral activity of DFMA was also studied by Del Sarto et al., (2020), demonstrating that the nucleoside analog DFMA was able to inhibit the viral replication and neuro-inflammation induced by ZKV *in vivo,* in addition to reducing lethality in A129 mice (28). Until then, DFMA had only been tested against flaviviruses, which generally have a conserved RdRp enzyme among them, considered an excellent target for antiviral development (36, 37). Our hypothesis was that DFMA could potentially exert a similar effect against members of the *Coronaviridae* family, given that both are positive single-stranded RNA viruses.

Considering the importance of assessing the antiviral capability not only for SARS-CoV-2, but against other coronaviruses, given the fact that the *Coronaviridae* family has already presented several members with a wide capacity to cause serious impacts on our society, the potential antiviral activity of DFMA was evaluated for MHV (Murine hepatitis virus). MHV, like SARS-CoV and SARS-CoV-2, is a betacoronavirus that infects mice naturally, capable of causing hepatitis, which can evolve into lethality of these animals. (23). Working with the MHV requires laboratories and animal facilities BSL-2, unlike SARS-CoV-2 virus, which requires BSL-3 laboratories. Therefore, as demonstrated by our group, the MHV-3 infection model, when inoculated intranasally, effectively induces lung damage (24). The model makes it possible to evaluate various parameters within BSL-2 laboratories, thus being used for screening and validation of potential antivirals with promising *in vitro* results. Our data showed DFMA as a promise putative antiviral, as it decreased the titers of both MHV-A59 and MHV-3, reducing virus quantification in the supernatant of infected L929 cells starting from 10 µM, similar to shown against ZKV as demonstrated by Del Sarto et al. where DFMA also reduced viral titers concentrations starting at 10 µM (28). DFMA showed excellent *in vitro* results against SARS-CoV-2, with a modest selectivity index of 6.22, which led us to take the next step towards *in vivo* studies.

Upon transitioning to *in vivo* experiments, using a treatment protocol similar to that replicated in other studies (28, 29), DFMA significantly reduced viral load in the lungs of mice across all three infection models MHV-3, MHV-A59, and SARS-CoV-2 as measured by qPCR. In addition, the results showed that the reduction in viral load likely resulted in effects on the production of different inflammatory mediators when quantified in the lungs of all infection models tested, as well as in the serum of mice in the experimental model of severe COVID using MHV-3. Like DFMA, the reduction of inflammatory mediators in the lungs of K18-hACE2 mice infected with SARS-CoV-2 was previously reported with the administration of molnupiravir, an antiviral molecule already approved as a drug against SARS-CoV-2 (38).

Systemic data demonstrate that, in a severe COVID model, although DFMA reduced viral load and inflammatory mediators, it did not significantly reduce the granulocyte/lymphocyte ratio in the MHV-3 model. However, in the mild COVID model, DFMA significantly reduced the granulocyte/lymphocyte ratio. The fact that there were no differences in the more severe COVID model raises the possibility that in the future, DFMA in synergy with some pro-resolutive drug may further enhance the prognostic and systemic effects, given that exacerbated inflammation and cytokine storm are also considered key factors in the evolution of COVID-19 patients.

In conclusion, DFMA showed promising results regarding its antiviral action both *in vitro* and *in vivo*, for MHV and SARS-CoV-2. Additionally, the fact that different members of the *Coronaviridae* family were tested suggests that DFMA may be an excellent candidate against future coronaviruses that may emerge in the coming years. Further studies need to be conducted to confirm the action and possible use of DFMA in humans. The present work demonstrates the value of using the MHV-3 and MHV-A59 as platforms for screening potential antiviral drugs against coronaviruses, depending on the mechanism of action.

### Financial support

This work was financially supported by the National Institute of Science and Technology in Dengue and Host-Microorganism Interactions (INCT Dengue), funded by the Brazilian National Science Council (CNPq, Brazil, grant number 465425/2014-3) and the Minas Gerais Foundation for Science (FAPEMIG, Brazil, grant number 25036/2014-3). Additional funding was provided by the Rede de Pesquisa em Imunobiológicos e Biofármacos para Terapias Avançadas e Inovadoras (ImunoBioFar) program, supported by FAPEMIG (grant numbers RED-00202-22, 29568-1, APQ-02281-18, APQ-02618-23, APQ-04650-23, and APQ-04983-24). This study also received partial funding from the Coordination for the Improvement of Higher Education Personnel (CAPES, Brazil, grant number 88881.507175/2020-01) and FINEP (Financier of Studies and Projects) under the MCTI/FINEP–MS/SCTIE/DGITIS/CGITS program (grant number 6205283B-BB28-4F9C-AA65-808FE4450542). Additional support was provided by the NIH (grant AI-RO1-161570) and Emory University’s CFAR (NIH grant P30 AI050409), both awarded to RFS.

## Acknowledgements

We extend our gratitude to Ilma Marçal de Souza, Rosemeire Oliveira, Letícia Soldati, and Tânia Colina for their invaluable technical assistance. We also acknowledge the support of the Technological Center for Advanced and Innovative Therapies (CT-Terapias) at UFMG. VVC and ASM express their gratitude for the Para Mulheres na Ciência Prize, awarded by L’Oréal, UNESCO, and the Brazilian Academy of Sciences (ABC). We thank the Laboratório Federal de Defesa Agropecuária (LFDA-MG), Ministry of Agriculture, Livestock, and Food Supply (MAPA), for enabling and supporting our activities in the BSL-4 laboratory. Finally, we acknowledge the Animal Biosafety Level 3 Laboratory at UFMG (Laboratório Institucional de Pesquisa, LIPq), the Multiuser Laboratory Center (CELAM), and the Biosafety Level 3 Laboratory (NB3-ICB) for their essential contributions.

## References

(1) Bull-Otterson, L., Saydah, S. B. S., Boehmer, T. K., Adjei, S., Gray, S., & Harris, A. M. (2022). Post-COVID Conditions Among Adult COVID-19 Survivors Aged 18–64 and ≥65 Years. Morbidity and Mortality Weekly Report Post-COVID, 71(21), 713–717. 10.15585/mmwr.mm7121e1

(2) Davis, H. E., McCorkell, L., Vogel, J. M., & Topol, E. J. (2023). Long COVID: major findings, mechanisms and recommendations. Nature Reviews Microbiology, 21(3), 133–146. 10.1038/s41579-022-00846-2

(3) Littlefield, K. M., Watson, R. O., Schneider, J. M., Neff, C. P., Yamada, E., Zhang, M., Campbell, T. B., Falta, M. T., Jolley, S. E., Fontenot, A. P., & Palmer, B. E. (2022). SARS-CoV-2-specific T cells associate with inflammation and reduced lung function in pulmonary post-acute sequalae of SARS-CoV-2. PLoS Pathogens, 18(5), 1–20. 10.1371/journal.ppat.1010359

(4) Monje, M., & Iwasaki, A. (2022). The neurobiology of long COVID. Cell Press, 110, 19–21. 10.1016/j.neuron.2022.10.006 SUMMARY

(5) Xie, Y., Xu, E., Bowe, B., & Al-Aly, Z. (2022). Long-term cardiovascular outcomes of COVID-19. Nature Medicine, 28(3), 583–590. 10.1038/s41591-022-01689-3

(6) Silva Andrade B, Siqueira S, de Assis Soares WR, de Souza Rangel F, Santos NO, dos Santos Freitas A, Ribeiro da Silveira P, Tiwari S, Alzahrani KJ, Góes-Neto A, et al. Long-COVID and Post-COVID Health Complications: An Up-to-Date Review on Clinical Conditions and Their Possible Molecular Mechanisms. Viruses. 2021; 13(4):700. 10.3390/v13040700

(7) Nicola, M., Alsafi, Z., Sohrabi, C., Kerwan, A., & Al-jabir, A. (2020). The socio-economic implications of the coronavirus pandemic (COVID-19): International Journal of Surgery, 78 (2020) 185–193. 10.1016/j.ijsu.2020.04.018

(8) Huang, C., Wang, Y., Li, X., Ren, L., Zhao, J., Hu, Y., Zhang, L., Fan, G., Xu, J., Gu, X., Cheng, Z., Yu, T., Xia, J., Wei, Y., Wu, W., Xie, X., Yin, W., Li, H., Liu, M., … Cao, B. (2020). Clinical features of patients infected with 2019 novel coronavirus in Wuhan, China. The Lancet, 395(10223), 497–506. 10.1016/S0140-6736(20)30183-5

(9) Montazersaheb, S., Hosseiniyan Khatibi, S. M., Hejazi, M. S., Tarhriz, V., Farjami, A., Ghasemian Sorbeni, F., Farahzadi, R., & Ghasemnejad, T. (2022). COVID-19 infection: an overview on cytokine storm and related interventions. Virology Journal, 19(1), 1–15. 10.1186/s12985-022-01814-1

(10) Cromer, D., Steain, M., Reynaldi, A., Schlub, T. E., Wheatley, A. K., Juno, J. A., Kent, S. J., Triccas, J. A., Khoury, D. S., & Davenport, M. P. (2022). Neutralising antibody titres as predictors of protection against SARS-CoV-2 variants and the impact of boosting: a meta-analysis. Lancet Microbe 2022;, *January*, e52–e61. 10.1016/S2666-5247(21)00267-6

(11) Li, R., Liu, J., & Zhang, H. (2021). The challenge of emerging SARS-CoV-2 mutants to vaccine development. Journal of Genetics and Genomics, 48(2), 102–106. 10.1016/j.jgg.2021.03.001

(12) Rössler A, Riepler L, Bante D, von Laer D, Kimpel J. SARS-CoV-2 Omicron Variant Neutralization in Serum from Vaccinated and Convalescent Persons. N Engl J Med. 2022 Feb 17;386(7):698–700. doi: 10.1056/NEJMc2119236. Epub 2022 Jan 12. PMID: 35021005; PMCID: PMC8781314.

(13) Viana, R., Moyo, S., Amoako, D. G., Tegally, H., Scheepers, C., Althaus, C. L., Anyaneji, U. J., Bester, P. A., Boni, M. F., Chand, M., Choga, W. T., Colquhoun, R., Davids, M., Deforche, K., Doolabh, D., du Plessis, L., Engelbrecht, S., Everatt, J., Giandhari, J., … de Oliveira, T. (2022). Rapid epidemic expansion of the SARS-CoV-2 Omicron variant in southern Africa. Nature, 603(7902), 679–686. 10.1038/s41586-022-04411-y

(14) Wang, P., Nair, M. S., Liu, L., Iketani, S., Luo, Y., Guo, Y., Wang, M., Yu, J., Zhang, B., Kwong, P. D., Graham, B. S., Mascola, J. R., Chang, J. Y., Yin, M. T., Sobieszczyk, M., Kyratsous, C. A., Shapiro, L., Sheng, Z., Huang, Y., & Ho, D. D. (2021). Antibody resistance of SARS-CoV-2 variants B.1.351 and B.1.1.7. Nature, 593(7857), 130–135. 10.1038/s41586-021-03398-2

(15) Channappanavar, R., & Perlman, S. (2017). Pathogenic human coronavirus infections: causes and consequences of cytokine storm and immunopathology. Seminars in Immunopathology, 39(5), 529–539. 10.1007/s00281-017-0629-x

(16) Bauer, R. N., Teterina, A., Shivram, H., McBride, J., Rosenberger, C. M., Cai, F., Bao, M., Tsai, L., Gordon, O., Lee, I. T., Wallin, J. J., Porter, D., Juneja, K., Camus, G., Rosas, I. O., & Wildum, S. (2023). Prognostic value of severe acute respiratory syndrome coronavirus-2 viral load and antibodies in patients hospitalized with COVID-19. Clinical and Translational Science, 16(6), 1049–1062. 10.1111/cts.13511

(17) Owen, D. R., Allerton, C. M. N., Anderson, A. S., Aschenbrenner, L., Avery, M., Berritt, S., Boras, B., Cardin, R. D., Carlo, A., Coffman, K. J., Dantonio, A., Di, L., Eng, H., Ferre, R. A., Gajiwala, K. S., Gibson, S. A., Greasley, S. E., Hurst, B. L., Kadar, E. P., … Zhu, Y. (2021). An oral SARS-CoV-2 Mpro inhibitor clinical candidate for the treatment of COVID-19. Science, 374(6575), 1586–1593. 10.1126/science.abl4784

(18) Brown, A. J., Won, J. J., Graham, R. L., Dinnon, K. H., Sims, A. C., Feng, J. Y., Cihlar, T., Denison, M. R., Baric, R. S., & Sheahan, T. P. (2019). Broad spectrum antiviral remdesivir inhibits human endemic and zoonotic deltacoronaviruses with a highly divergent RNA dependent RNA polymerase. Antiviral Research Journal, 169(January).

(19) Wang Y, Zhang D, Du G, Du R, Zhao J, Jin Y, Fu S, Gao L, Cheng Z, Lu Q, Hu Y, Luo G, Wang K, Lu Y, Li H, Wang S, Ruan S, Yang C, Mei C, Wang Y, Ding D, Wu F, Tang X, Ye X, Ye Y, Liu B, Yang J, Yin W, Wang A, Fan G, Zhou F, Liu Z, Gu X, Xu J, Shang L, Zhang Y, Cao L, Guo T, Wan Y, Qin H, Jiang Y, Jaki T, Hayden FG, Horby PW, Cao B, Wang C. Remdesivir in adults with severe COVID-19: a randomised, double-blind, placebo-controlled, multicentre trial. Lancet. 2020 May 16;395(10236):1569–1578. doi: 10.1016/S0140-6736(20)31022-9. Epub 2020 Apr 29. Erratum in: Lancet. 2020 May 30;395(10238):1694. doi: 10.1016/S0140-6736(20)31204-6.

(20) Mahase, E. (2021). Covid-19: Molnupiravir reduces risk of hospital admission or death by 50% in patients at risk, MSD reports. BMJ (Clinical Research Ed.), 375(October), n2422. 10.1136/bmj.n2422

(21) MERK. (2021). Merck-and-Ridgebacks-Investigational-Oral-Antiviral-Molnupiravir-Reduced-the-Risk-of-Hospitalization-or-Death-by-Approximately-50-Per-C1CSM. 1–6.

(22) Urakova, N., Kuznetsova, V., Crossman, D. K., Sokratian, A., Guthrie, D. B., Kolykhalov, A. A., Lockwood, M. A., Natchus, M. G., Crowley, M. R., Painter, G. R., Frolova, E. I., & Frolov, I. (2018). β- d - N 4 -Hydroxycytidine Is a Potent Anti-alphavirus Compound That Induces a High Level of Mutations in the Viral Genome. Journal of Virology, 92(3), 1–22. 10.1128/jvi.01965-17

(23) Lucchiari MA, Pereira CA, Kuhn L, Lefkovits I. The pattern of proteins synthesized in the liver is profoundly modified upon infection of susceptible mice with mouse hepatitis virus 3. Res Virol. 1992 Jul-Aug;143(4):231-40. doi: 10.1016/s0923-2516(06)80111-1. PMID: 1329165; PMCID: PMC7135047.

(24) Andrade, A. C. dos S. P., Campolina-Silva, G. H., Queiroz-Junior, C. M., Oliveira, L. C. de, Lacerda, L. de S. B., Gaggino, J. C. P., Souza, F. R. O. de, Chaves, I. de M., Passos, I. B., Teixeira, D. C., Bittencourt-Silva, P. G., Valadão, P. A. C., Rossi-Oliveira, L., Antunes, M. M., Figueiredo, A. F. A., Wnuk, N. T., Jairo R. Temerozo, E., Ferreira, A. C., Cramer, A., … Costaa, V. V. (2021). PATHOGENESIS AND IMMUNITY A biosafety level 2 ouse model for studying Betacoronavirus-Induced Acute Lung Damage and Systemic Manifestations. September, 1–18.. 10.1128/JVI.01276-21.

(25) Campolina-Silva, G., Andrade, A. C. dos S. P., Couto, M., Bittencourt-Silva, P. G., Queiroz-Junior, C. M., Lacerda, L. de S. B., Chaves, I. de M., de Oliveira, L. C., Marim, F. M., Oliveira, C. A., da Silva, G. S. F., Teixeira, M. M., & Costa, V. V. (2023). Dietary Vitamin D Mitigates Coronavirus-Induced Lung Inflammation and Damage in Mice. Viruses, 15(12). 10.3390/v15122434

(26) Pimenta, J. C., Beltrami, V. A., Oliveira, B. D. S., Queiroz-Junior, C. M., Barsalini, J., Teixeira, D. C., Souza-Costa, L. P. de, Lima, A. L. D., Machado, C. A., Parreira, B. Z. S. G., Santos, F. R. da S., Costa, P. A. C., Lacerda, L. D. S. B., Gonçalves, M. R., Chaves, I. de M., Couto, M. G. G. Do, Costa, V. R. de M., Nóbrega, N. R. C., Silva, B. L., … Costa, V. V. (2024). Neuropsychiatric sequelae in an experimental model of post-COVID syndrome in mice. BioRxiv, 2024.01.10.575003. 10.1101/2024.01.10.575003.

(27) Guimaraes LC, Costa PAC, Scalzo Júnior SRA, Ferreira HAS, Braga ACS, de Oliveira LC, Figueiredo MM, Shepherd S, Hamilton A, Queiroz-Junior CM, da Silva WN, da Silva NJA, Rodrigues Alves MT, Santos AK, de Faria KKS, Marim FM, Fukumasu H, Birbrair A, Teixeira-Carvalho A, de Aguiar RS, Mitchell MJ, Teixeira MM, Vasconcelos Costa V, Frezard F, Guimaraes PPG. Nanoparticle-based DNA vaccine protects against SARS-CoV-2 variants in female preclinical models. Nat Commun. 2024 Jan 18;15(1):590. doi: 10.1038/s41467-024-44830-1. PMID: 38238326; PMCID: PMC10796936.

(28) Del Sarto, J. L., Rocha, R. de P. F., Bassit, L., Olmo, I. G., Valiate, B., Queiroz-Junior, C. M., Pedrosa, C. da S. G., Ribeiro, F. M., Guimarães, M. Z., Rehen, S., Amblard, F., Zhou, L., Cox, B. D., Gavegnano, C., Costa, V. V., Schinazi, R. F., & Teixeira, M. M. (2020). 7-Deaza-7-fluoro-2′-C-methyladenosine inhibits Zika virus infection and viral-induced neuroinflammation. Antiviral Research, 180(June), 1–9. 10.1016/j.antiviral.2020.104855.

(29) Zandi, K., Bassit, L., Amblard, F., Cox, B. D., Hassandarvish, P., Moghaddam, E., Yueh, A., Rodrigues, G. O. L., Passos, I., Costa, V. V., AbuBakar, S., Zhou, L., Kohler, J., Teixeira, M. M., & Schinazi, R. F. (2019). Nucleoside analogs with selective antiviral activity against dengue fever and Japanese encephalitis viruses. Antimicrobial Agents and Chemotherapy, 63(7), 1–15. 10.1128/AAC.00397-19

(30) Chan, A. S., & Rout, A. (2020). Use of Neutrophil-to-Lymphocyte and Platelet-to Lymphocyte Ratios in COVID-19. Journal of Clinical Medicine Research, 12(7), 448–453. 10.14740/jocmr4240

(31) Li, X., Liu, C., Mao, Z., Xiao, M., Wang, L., Qi, S., & Zhou, F. (2020). Predictive values of neutrophil-to-lymphocyte ratio on disease severity and mortality in COVID-19 patients: a systematic review and meta-analysis. Critical Care, 24(1), 1–10. 10.1186/s13054-020-03374-8

(32) Su, S., Wong, G., Shi, W., Liu, J., Lai, A. C. K., Zhou, J., Liu, W., Bi, Y., & Gao, G. F. (2016). Epidemiology, Genetic Recombination, and Pathogenesis of Coronaviruses. Trends in Microbiology, 24(6), 490–502. 10.1016/j.tim.2016.03.003

(33) Qiu, W., Chu, C., Mao, A., & Wu, J. (2018). The impacts on health, society, and economy of SARS and H7N9 Outbreaks in China: A Case Comparison Study. Journal of Environmental and Public Health, 2018. 10.1155/2018/2710185

(34) Zheng C, Shao W, Chen X, Zhang B, Wang G, Zhang W. Real-world effectiveness of COVID-19 vaccines: a literature review and meta-analysis. Int J Infect Dis. 2022 Jan;114:252–260. doi: 10.1016/j.ijid.2021.11.009. Epub 2021 Nov 17. PMID: 34800687; PMCID: PMC8595975.

(35) Bhattacharya, M., Chatterjee, S., Sharma, A. R., Agoramoorthy, G., & Chakraborty, C. (2021). D614G mutation and SARS-CoV-2: impact on S-protein structure, function, infectivity, and immunity. Applied Microbiology and Biotechnology, 105(24), 9035–9045. 10.1007/s00253-021-11676-2

(36) Lim, S. P., Noble, C. G., & Shi, P. Y. (2015). The dengue virus NS5 protein as a target for drug discovery. Antiviral Research, 119(April), 57–67. 10.1016/j.antiviral.2015.04.010.

(37) Rawlinson, S., Pryor, M., Wright, P., & Jans, D. (2006). Dengue Virus RNA Polymerase NS5: A Potential Therapeutic Target? Current Drug Targets, 7(12), 1623– 1638. 10.2174/138945006779025383

(38) Papinia, C., Ullah, I., Amalendu P. Ranjan, Zhang, S., Wu, Q., Spasov, K. A., Zhang, C., Mothes, W., Crawford, J. M., Lindenbach, B. D., D. Uchil, P., Kumar, P., Jorgensen, W. L., & S. Anderson, K. (2024). Proof- of- concept studies with a computationally designed Mpro inhibitor as a synergistic combination regimen alternative to Paxlovid. 121, 10. 10.1073/pnas.2320713121.

